# An Aged Microenvironment Increases CAR T Cell Cytotoxicity but Impairs Therapeutic Efficacy

**DOI:** 10.1101/2025.05.04.652106

**Authors:** Mona Kabha, Noy Rosenzweig, Akmaral Rakhymzhanova, Orna Atar, Cyrille J. Cohen, Noga Ron-Harel

**Affiliations:** Faculty of Biology, Technion-Israel Institute of Technology, Haifa, Israel; The Laboratory of Tumor Immunology and Immunotherapy, The Mina and Edvard Goodman Faculty of Life Sciences, Bar-Ilan University, Israel

## Abstract

**Background:** Cancer disproportionately affects the elderly, who are often less able to tolerate traditional cytotoxic therapies, and may benefit from T cell–based immunotherapies. However, studies evaluating the efficacy of T cell immunotherapy in aged mice are limited and yield inconsistent results, while clinical data are largely retrospective.

**Methods:** Here, we used a murine model of Chimeric Antigen Receptor (CAR) T cell therapy to investigate how aging influences efficacy, from CAR T cell production to in vivo anti-tumor activity.

**Results:** We found that aging reduced CAR T cell production yields and altered their phenotype and function. Aged CAR T cells were predominantly effector memory CD4⁺ T cells, whereas young CAR T cells were primarily central memory CD8⁺ T cells. Functionally, aged CAR T cells exhibited non-specific cytotoxicity, driven by constitutive degranulation and elevated granzyme B secretion independent of CAR expression. This phenotype was induced by the aged microenvironment, as young T cells transferred into aged hosts adopted similar behavior. In vivo, young CAR T cells efficiently reduced tumor burden in young leukemia-bearing hosts, but were not effective in aged hosts, where the aged microenvironment impaired CAR T cell persistence.

**Conclusion:** These findings indicate that aging impacts CAR T cell therapy at multiple levels, from manufacturing to therapeutic efficacy, highlighting the need to design tailored immunotherapies for elderly patients.

## Background

Cancer disproportionally affects elderly people, and aged patients are less able to tolerate the detrimental side effects of classical therapies, such as systemic chemotherapy and irradiation, which cause excessive damage to healthy tissues. Hence, aged patients could greatly benefit from more specific therapeutic approaches such as immunotherapy based on endogenous or genetically engineered T cells to directly target tumor cells. The choice of cancer immunotherapy must therefore consider age-related changes in T cell immunity^1^.

Chimeric antigen receptor (CAR) T cells are genetically engineered T cells designed to recognize and attack tumor-specific antigens^2^. These synthetic receptors consist of an extracellular antigen-binding domain, typically a single-chain variable fragment (scFv) derived from an antibody, linked to intracellular signaling domains, including the CD3ζ activation domain and one or more co-stimulatory domains such as CD28 or 4-1BB. Upon engagement with their target antigen, CARs initiate intracellular signaling cascades that lead to T cell activation, proliferation, and acquisition of effector functions. Activated CAR T cells eliminate target cells through the release of cytotoxic granules (e.g., perforin and granzyme B), as well as proinflammatory cytokines and chemokines^3^.

The expression on CAR T cells of exhaustion markers such as PD-1, Tim-3, and LAG-3, which are also elevated in aged T cells, has been associated with poor clinical outcomes^4,5^. However, to date, no clinical studies have directly examined the impact of T cell aging on CAR T cell immunotherapy outcomes^6^. Retrospective analyses of the Tisagenlecleucel (JULIET trial)^7^ and Axicabtagene Ciloleucel (ZUMA-1 trial)^8^ found that age did not significantly affect the efficacy of anti-CD19 CAR T-cell therapies in relapsed or refractory B-cell malignancies. Both trials included patients up to 76 years of age, and reported no significant differences in overall or complete response rates between those over 65 and younger patients. Similarly, post-authorization studies for relapsed or refractory multiple myeloma (RR MM)^9^ showed comparable outcomes between younger and older patients. These findings suggest that aging does not compromise CAR T-cell efficacy; however, older patients had a higher risk of severe adverse events, including cytokine release syndrome (CRS) and neurological toxicity. In addition, aging increases the likelihood of CAR-T product manufacturing failure^10^, raising concerns that retrospective analyses may introduce selection bias by including only older patients whose T cells were fit enough to successfully complete the CAR T-cell production process^6^.

Preclinical studies in animal models evaluating the impact of aging on immunotherapy efficacy have yielded inconsistent results. Some studies reported enhanced cytotoxicity in CD8^+^ T cells from aged mice, attributed to increased secretion of perforin and granzyme B (GzB)^11,12^. In contrast, other studies indicated that CAR T cells derived from aged donor mice exhibit functional impairments compared to those from young donors, including lower transduction efficiency, reduced expansion, and diminished levels of key signaling molecules such as phosphorylated ERK, Akt, Stat3, and Stat5, ultimately leading to reduced cytotoxicity^13^.

In this study, we utilized the well-established CD19 CAR T cell model^14^, and demonstrated that aging negatively impacted CAR T cell production, phenotype, and function. Aged CAR T cells were predominantly comprised of effector memory CD4^+^ T cells, whereas young CAR T cells primarily consisted of central memory CD8^+^ T cells. Functionally, aged CAR T cells exhibited non-specific cytotoxicity, targeting and killing cells independently of CAR expression or direct contact. This excessive killing was driven by continuous degranulation and elevated GzB secretion. The aged microenvironment played a key role in inducing this phenotype, as young T cells transferred into aged hosts displayed a similar phenotype. In studies employing the ML21 leukemia model^15^, young CAR T cells effectively reduced tumor burden in young leukemia-bearing hosts but were less effective in aged hosts. Within the aged microenvironment, CAR T cells exhibited reduced persistence and were less efficient in reducing tumor burden.

## Methods

### Mice

Young (8-12 week old) and aged (20-23 month old) C57BL/6JOlaHsd female mice were purchased from Envigo (Israel). For aging experiments, 8 months old retired breeders were maintained for additional 12-15 months. Transgenic C57Bl/6 Rosa26td^Tomato/+^ OTII mice were a gift from Prof. Ziv Shulman (The Weizmann Institute of Science). All mice were housed under specific pathogen-free conditions at The Technion Pre–Clinical Research Authority and used in accordance with animal care guidelines of the Institutional Animal Care and Use Committee.

### Cell Lines

The A20 cell line was cultured in RPMI 1640 medium supplemented with 10% FBS (Gibco), 1% Penicillin/Streptomycin (Gibco), 1% HEPES buffer, and 50μM β-mercaptoethanol. HEK293T cells (Human embryonic kidney 293 cells) were cultured in Dulbecco’s modified Eagle’s medium (DMEM) supplemented with 10% of FBS and 1% Penicillin/Streptomycin.The cultures were maintained at 37℃ with 5% CO2 and split every 3-4 days.

### Construction of retroviral vectors encoding CARs

Second-generation CAR constructs were prepared by cloning the variable region of the 1D3 hybridoma that specifically recognizes murine CD19 (mCD19 scFv) into the retroviral pBullet vector, including a gene encoding the GFP label (A gift from Prof. Zelig Eshchar’s lab). The scFv was connected via the carboxyl terminus of the heavy chain variable domain VH region to a myc tag, a short peptide sequence used to facilitate the detection of cells that expressed the construct, CD8 hinge, CD8 transmembrane domain, and CD28-CD3ζ intracellular domains, and cloned into the retroviral pBullet vector, followed by an internal ribosome entry site (IRES) and GFP.

### Generating CAR T cells from aged and young mice

HEK293T cells were transfected with CAR retroviral vectors and Peco packaging plasmids, using the CalFectin reagent. Virus-containing supernatants were harvested after 48 hours of culture and used for transduction.

Retroviral transduction of primary mouse T cells: Splenocytes were harvested from young (8-12 weeks old) and aged (20-22 months old) C57Bl/6 mice, and T cells were isolated by magnetic separation using EasySep™ Mouse T Cell Isolation Kit (STEMCELL Technologies, catalog #19851) according to the manufacturer’s instructions. The isolated cells were resuspended in stimulation media (complete RPMI medium supplemented with 1% non-essential amino acids, 1% Sodium Pyruvate Solution, and 100 U/ml recombinant murine IL-2 (Peprotech, catalog #212-12-50)), stimulated either using weak (0.5 μg/ml α-CD28; BioXcell BE0015-1 and 1 μg/ml α-CD3; BioXcell BE0001-1) or strong (4 μg/ml α-CD28 and 1 μg/ml α-CD3) activation, and infected with a retrovirus, using retronectin (TaKara, Catalog #T100A) according to the manufacturer’s instructions. CAR T cells were cultured at 37 °C and 5 % CO2 in cRPMI supplemented with IL-2 (100U/ml).

### CAR T cell sorting

GFP^+^ cells were sorted using a FACS ARIA–IIIU (BD Biosciences) or Bigfoot (Thermo Fisher Scientific) Cell Sorters. After sorting, cells were cultured in complete RPMI medium supplemented with recombinant murine IL-2 (100U/ml).

### Killing assays

Coculture: A20 cells were labeled with CellTrace Violet (CTV) (Thermo Fisher Scientific, Catalog# C34557) following the manufacturer’s protocol, and co-cultured with young or aged control T or CAR T cells at varying target-to-effector (T:E) ratios. A20 cell death was assessed using either live cell imaging or flow cytometry. Transwell: Cytokine-mediated killing was measured using the Corning® HTS Transwell®-96 Permeable Support with a 0.4 µm Pore Polycarbonate Membrane (Corning, catalog #3381). Young and aged CAR T cells or control T cells were cultured in the upper chamber, while A20 cells were added to the lower chamber at a 1:5 T:E ratio for 10 hours. After incubation, the A20 cells were collected and stained with Zombie NIR™ Fixable Viability Kit (Biolegend, catalog # 423105) dye to assess target cell death.

### Flow cytometry

For cell surface staining, cells were resuspended in separation buffer (PBS containing 2% FBS and 2 mM EDTA) and incubated on ice for 20 minutes with antibody mixture, washed, and analyzed by flow cytometry. For intracellular staining, cells were activated using PMA and ionomycin (Biolegend, catalog # 423301) in the presence of Golgi Stop (Brefeldin A; Biolegend, catalog # 420601) for 5 hours, and stained using a True-Nuclear Transcription Factor Buffer Set (Biolegend, catalog # 424401), following the manufacturer’s protocol.

List of antibodies used:

| Antibody | Supplier | Catalog number |
| --- | --- | --- |
| PE anti-mouse CD4 | BioLegend | 100408 |
| APC anti-mouse CD8a | BioLegend | 100712 |
| PE anti-mouse CD69 | BioLegend | 104508 |
| Brilliant Violet 510™ anti-mouse/human CD44 | Biolegend | 103043 |
| Alexa Fluor® 700 anti-mouse CD62L | Biolegend | 104426 |
| eFluor™ 450 IFN gamma Monoclonal Antibody (XMG1.2) | ThermoFisher Scientific | 48-7311-82 |
| Brilliant Violet 510 anti-mouse CD107a (LAMP-1) | Biolegend | 121629 |
| Alexa Fluor® 647 anti-human/mouse Granzyme B Recombinant Antibody | Biolegend | 396421 |
| Propidium Iodide Solution | Biolegend | 421301 |
| APC anti-mouse CD19 Antibody | Biolegend | 152409 |
| Alexa Fluor® 647 anti-c-Myc Antibody | Biolegend | 626810 |
| CellTrace™ Violet Cell Proliferation Kit (CTV) | ThermoFisher Scientific | C34557 |
| Incucyte® Annexin V Orange Reagent for apoptosis | Sartorius | 4759 |
| Zombie NIR™ Fixable Viability Kit | Biolegend | 423105 |
| FITC anti-mouse CD8a | BioLegend | 100706 |

### Live cell imaging

Live cell imaging was performed by labeling target A20 cells with Cell Trace Violet and co-culturing them with either young or aged control T cells or CAR T cells at a 1:1 effector-to-target ratio. Annexin V antibodies were added to the culture medium to mark apoptotic cells, and the percentage of CTV/Annexin V double-positive A20 cells was quantified over 24 hours using the Zeiss Celldiscoverer 7. Analysis was performed using Zen software.

### ELISA

Supernatants collected from the killing experiments were used to quantify Granzyme B and IFNγ secretion. The levels of these cytokines were determined using the Mouse Granzyme B DuoSet ELISA (R&D Systems, catalog #DY1865-05) and IFNγ DuoSet ELISA (R&D Systems, catalog #DY485-05) according to the manufacturer’s instructions.

### Adoptive T cell transfer

Young T cells were isolated from spleens of young C57Bl/6 Rosa26td^Tomato/+^ (8-12 week old) by magnetic isolation (StemCell), and 5X10^6^ cells were transferred i.v. into wild type C57Bl/6 young or aged recipients.

### ML21 Leukemia model and CAR T cell treatment

Young or aged mice were injected intraperitoneally (IP) with 200 mg/Kg Cyclophosphamide monohydrate (Sigma, Catalog # C0768-1G). After 1 day, 1X10^6^ ML21 cells were administered intravenously (iv) to the mice. Drinking water was supplemented with 20 mg/ml doxycycline for the entire experiment duration. On day 10, the mice received i.v. injection of either 2X10^6^ control or CAR T cells, and blood samples were collected via tail vein puncture every few days to assess tumor burden and CAR T cell persistency.

## Results

### Suboptimal activation impairs the production of CAR T cells from aged donors

To directly investigate how aging affects the efficacy of CAR T cell therapy, we utilized the CD19 CAR T cell model, which has demonstrated efficacy in both preclinical and clinical studies against CD19^+^ B cell malignancies^16,17^. A second-generation CAR construct targeting murine CD19 (mCD19; supplemental figure 1A) was generated and cloned into the GFP-labeled retroviral pBullet vector. A Myc tag^18^ was included to facilitate the detection of CAR-expressing T cells. CAR T cells were prepared using standard protocols^19^ (figure 1A), and their functionality was validated by assessing their ability to kill A20 B cell lymphoma cells expressing CD19, demonstrating a specific, dose-dependent cytotoxicity (supplemental figure 1B).

**Figure 1:**
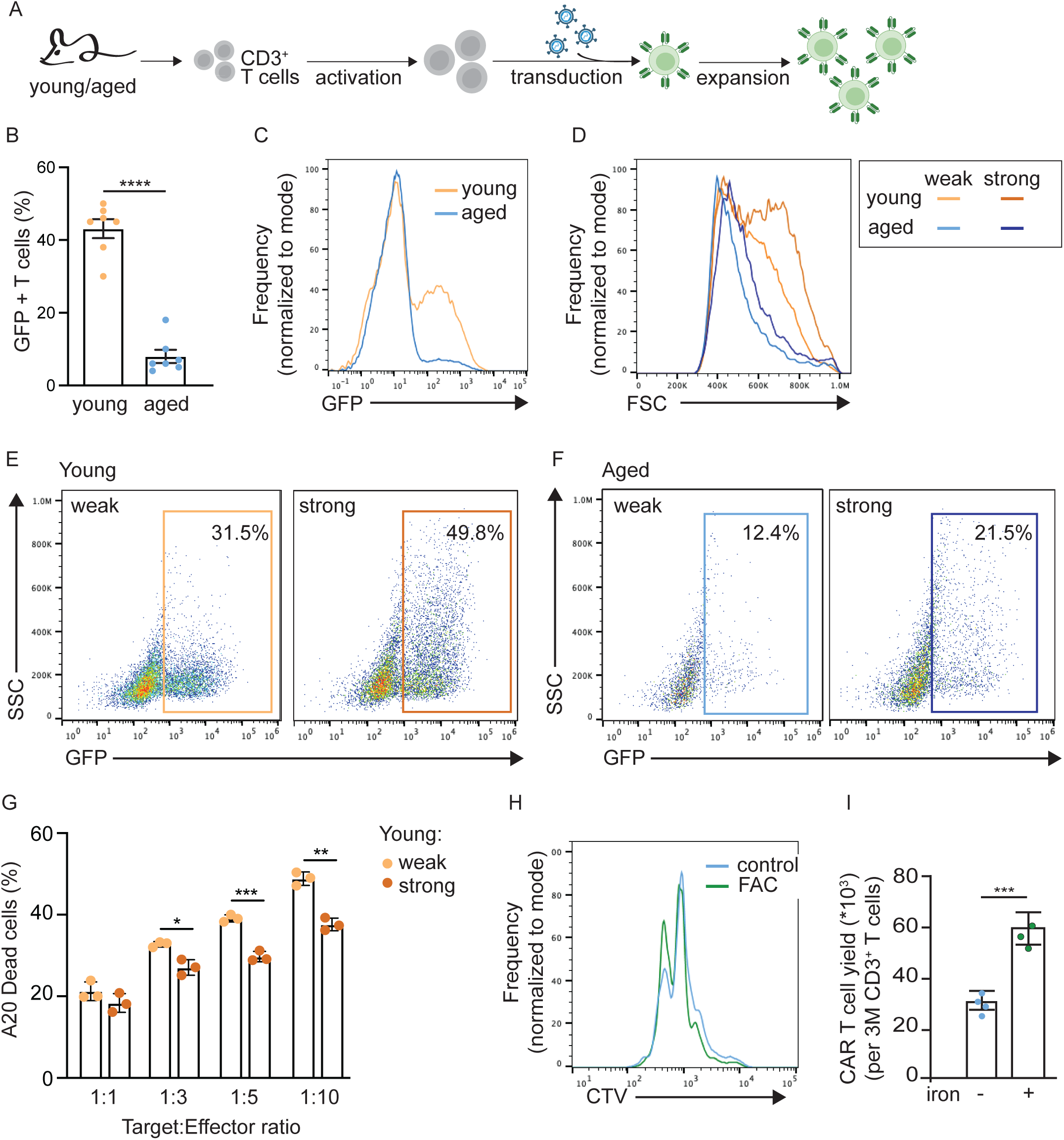
Suboptimal activation impairs the production of CAR T cells from aged donors. (A) Scheme showing experimental design. CD3^+^ T cells were isolated from the spleens of young or aged C57Bl/6 mice, activated, and transduced with retroviruses carrying the CAR plasmid. (B) Quantification of transduction efficiency by the percentage of GFP^+^ CAR T cells (each data point represent a pool of aged (n=5) or young (n=3) mice). (C) a representative FACS plot. (D) analysis of cell size, and (E-F) transduction efficiency in young versus aged T cells following a “weak” (0.5 μg/ml anti-CD28 and 1 μg/ml anti-CD3) or “strong” (4 μg/ml anti-CD28 and 1 μg/ml anti-CD3) stimulation. (G) A20 cells were labeled with CellTrace Violet (CTV) and co-cultured with young CAR T cells, prepared using either weak or strong activation at varying target-to-effector (T:E) ratios. A20 cell death was assessed using PI staining (data points are technical replicates of T cells pooled from 5 young mice). (H) Representative plot showing proliferation of aged CAR T cells prepared with or without supplementation with ferric ammonium citrate (FAC). (I) aged CAR T cell production yield with or without FAC supplementation (data points are technical replicates of T cells pooled from 6 aged mice). Bar graphs represent mean ± SEM (*p<0.05, **p<0.01, ***p<0.001, ****p<0.0001); one-way ANOVA with Tukey’s multiple comparisons test (G), or unpaired Student’s t test (B,I)).

Transduction efficiency in T cells from aged mice (20-23 months old) was below 10% and significantly lower than that in T cells from young mice (8-12 weeks old), which exhibited over 40% efficiency, as measured by GFP expression (figures 1B, 1C). To enable functional studies, we sought to enhance CAR T cell production from aged T cells. Since successful transduction requires pre-activation, standard CAR T cell protocols recommend low concentrations of anti-CD3 and anti-CD28 antibodies to minimize T cell exhaustion^20^. Under such “weak activation” conditions (0.5 μg/ml anti-CD28 and 1 μg/ml anti-CD3), fewer than 30% of aged T cells expressed the early activation marker CD69, compared to over 70% of young T cells (supplemental figure 1C). Increasing anti-CD28 concentrations slightly improved transduction efficiency (supplemental figure 1D), whereas increasing anti-CD3 had no effect (supplemental figure 1E). Thus, to improve CAR T cell production yield from aged cells, we employed a “strong activation” protocol (4 μg/ml anti-CD28 and 1 μg/ml anti-CD3), which significantly increased CD69 expression (supplemental figure 1F), promoted activation-induced cell growth (figure 1D), and improved transduction efficiency in both young and aged T cells (figures 1E, F). However, CAR T cells generated under this stronger activation were functionally inferior to those generated using the weak activation protocol (figure 1G). These findings prompted us to explore alternative strategies to improve CAR T cell production from aged donors.

Our previous studies demonstrated that T cells from aged mice fail to increase labile iron pools upon activation, and iron supplementation during activation enhances their proliferation^21^. Based on these findings, we tested whether iron supplementation during the initial activation step would improve aged CAR T cells production yield. GFP^+^ CAR T cell analysis revealed enhanced proliferation, as assessed by CellTrace Violet dye dilution, in CAR T cells generated using iron supplementation (figure 1H). This led to a significant increase in the yield of aged CAR T cells (figure 1I). This optimized protocol enabled the generation of sufficient numbers of aged CAR T cells to analyze their cellular composition and function in comparison to those from young donors.

### The cellular composition of CAR T cells is affected by donor age

CAR T cells are generated from bulk CD3^+^ T cells. During aging, T cell composition changes are manifested by a shrinkage of the naïve T cell compartment and accumulation of memory and terminally differentiated T cells^1^. To determine how donor age impacts CAR T cell phenotype, T cells derived from young and aged donors were analyzed at different steps during CAR T cell production (figure 2A). As expected, T cells isolated from young donors were predominantly CD4^+^ cells, with a naïve phenotype (CD62L^+^CD44^lo^; figure 2B), whereas the majority of T cells derived from aged donors were effector memory (EM) cells (CD62L^-^CD44^hi^; Figure 2C). After initial activation, young T cells acquired a central memory (CM) phenotype, showing an increase in the proportion of CD8^+^ T cells (figure 2D). Aged T cells maintained their EM phenotype and, like young T cells, exhibited an increased proportion of CD8^+^ cells (figure 2E). Successfully transduced, GFP^+^ CAR T from young donors were primarily CD8^+^ cells, with a CM phenotype (figure 2F). However, in T cells derived from aged donors, the majority of successfully transduced cells exhibited a CD4^+^ EM phenotype (figure 2G). Together, these data demonstrate that the final composition of CAR T cells is influenced by donor age, raising the important question of whether and how this might impact their functionality.

**Figure 2:**
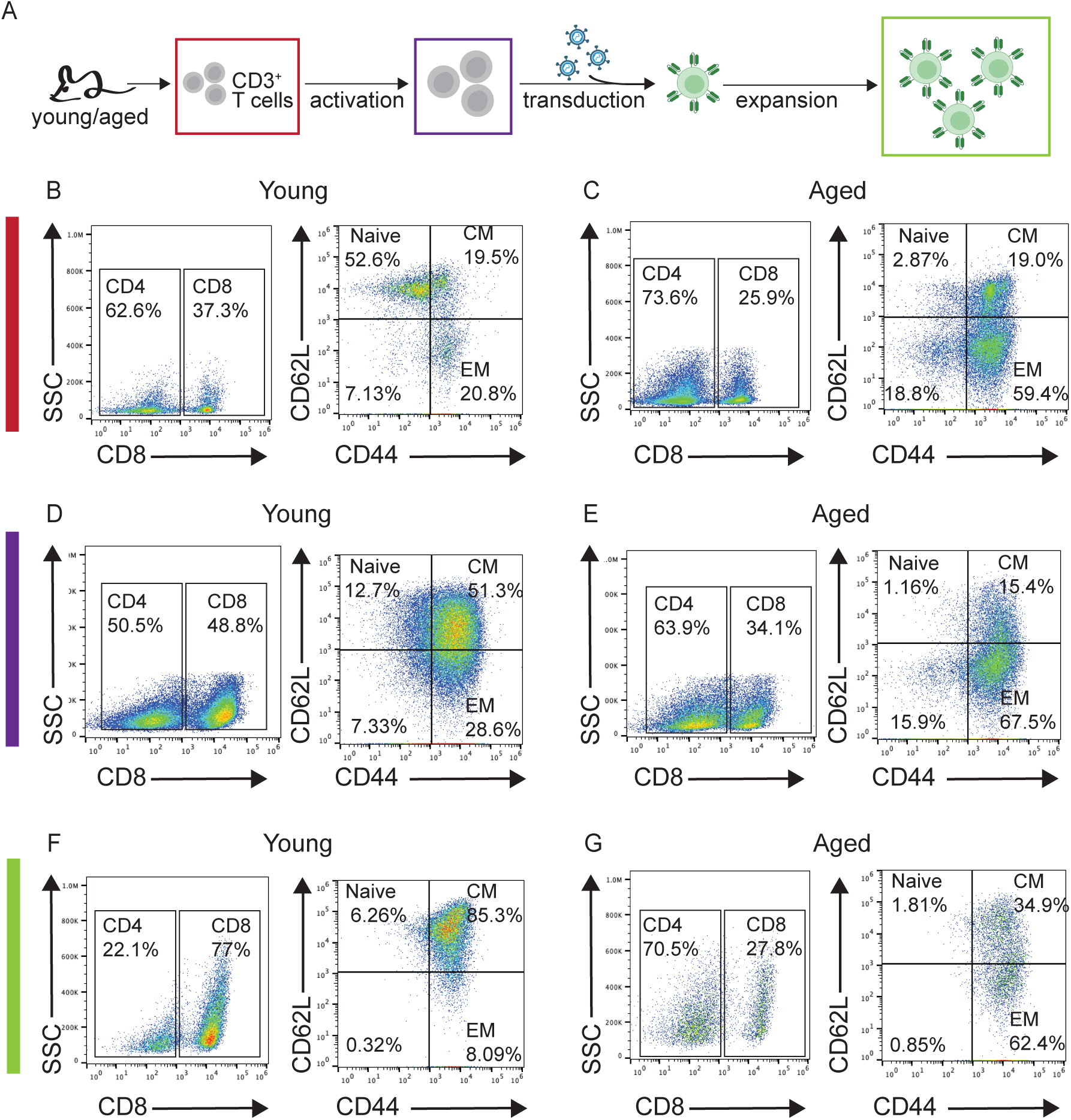
The cellular composition of CAR T cells is affected by donor age. (A) Experimental scheme. Young (n=3, pooled) and aged (n=5, pooled) T cells were activated, transduced with viruses carrying the CAR plasmid, and analyzed by flow cytometry at different steps of the production process, as indicated by the colored frames. (B,C) analysis of T cells immediately after isolation. (D,E) Analysis of T cells following initial activation, and (F,G) analysis gating on GFP^+^ CAR T cells. Panel shows representative data from 2 independent experiments.

### T cells from aged donors non-specifically kill target cancer cells independently of CAR expression

Upon binding to its target, CD19, the CAR triggers downstream signaling cascades that drive T cell activation, proliferation, and effector functions. Activated CAR T cells eliminate their targets through the secretion of cytotoxic granules containing perforin and GzB, proinflammatory cytokines (e.g., IFNγ), and chemokines^22^. To compare the functionality of young and aged CD19-CAR T cells, we monitored their killing efficiency against A20 target cells using live-cell imaging. A20 cells were stained with CellTrace Violet (CTV) and cocultured with either young or aged CAR T cells in media supplemented with anti-Annexin V antibodies to detect apoptosis. Cultures were observed for 24 hours, and the fraction of Annexin V^+^ A20 cells (CTV^+^) was quantified. Controls included A20 cells alone and A20 cells cocultured with primary young or aged T cells after pre-activation, without transduction.

Young CAR T cells efficiently killed their targets within the first hour of coculture, with 75% of A20 cells expressing Annexin V, compared to only 30% in the presence of control young T cells (figure 3A). In contrast, aged CAR T cells exhibited a delayed cytotoxic response, with a significant increase in Annexin V signal observed after 5 hours. Unexpectedly, aged control T cells killed target cells as efficiently as aged CAR T cells, suggesting that their cytotoxicity was independent of CAR expression (figure 3A).

**Figure 3:**
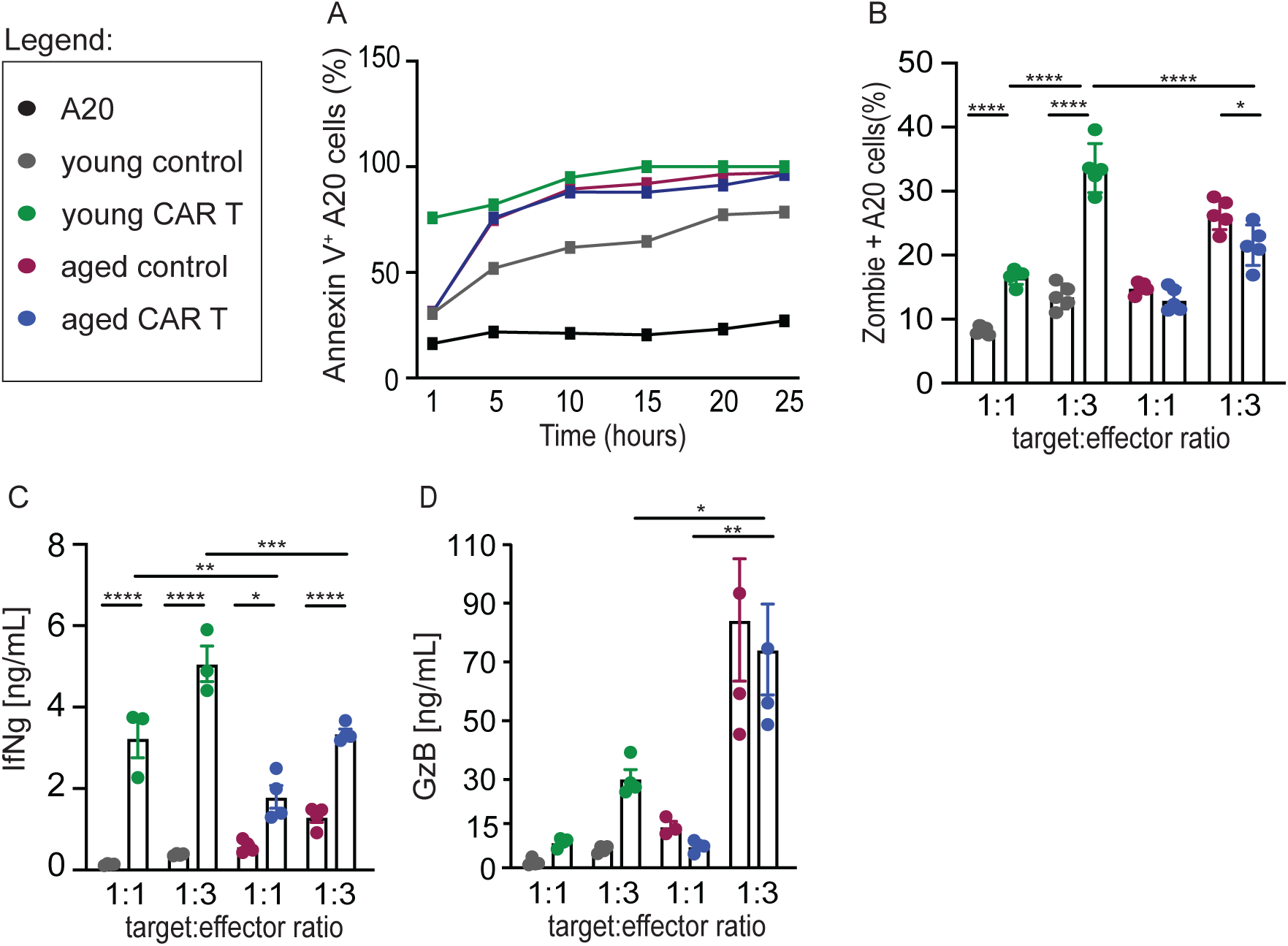
T cells from aged donors kill target cancer cells independently of CAR expression. (A) CAR T cells were produced from young (n=3, pooled) and aged (n=5, pooled) mice and cultured with CellTrace Violet-labeled A20 cells, at a 1:1 ratio. A20 killing was assessed over 24 hrs using live cell imaging and Annexin V staining. (B) Analysis of A20 killing after 10 hr coculture with young and aged CAR T cells at varying T: E ratios. A20 cell killing was evaluated by flow cytometry using Zombie Aqua staining. Media from this experiment was used to quantify IFNγ (C), and Granzyme B (GzB) (D) by ELISA. Bar graphs represent mean ± SEM. Data points are technical replicates. (*p<0.05, **p<0.01, ***p<0.001, ****p<0.0001); one-way ANOVA with Tukey’s multiple comparisons test.

To further validate these findings, we performed an additional killing assay at varying target-to-effector ratios, measuring target cell killing after 10 hours, the time point of maximal cytotoxicity (figure 3A). Young CAR T cells demonstrated specific, dose-dependent killing and were significantly more cytotoxic than aged CAR T cells, particularly at higher target-to-effector ratios. Notably, aged control T cells exhibited cytotoxicity comparable to that of aged CAR T cells (figure 3B).

Both young and aged CAR T cells responded to target recognition by secreting IFNγ. As expected, cultures with higher effector cell concentrations contained higher levels of IFNγ. Notably, young CAR T cells secreted more IFNγ than aged CAR T cells (figure 3C). GzB secretion by young CAR T cells increased as expected compared to young control cells (p<0.0001, one-way ANOVA; comparing only young control and CAR T cells at 1:3 T:E ratio; figure 3D). However, aged T cells secreted nearly twice as much GzB compared to young T cells, independent of target recognition (figure 3D). These findings suggest that T cell cytotoxicity in aged cells is at least partially mediated by continuous GzB secretion, raising the question of whether aged T cells possess inherent cytotoxicity independent of the ex vivo CAR T cell production process.

### Aged T cells eliminate target cells through the continuous secretion of GzB and IFNγ

To assess cytotoxicity mediated by aged T cells, T cells were purified from the spleens of young and aged mice and subjected to a brief chemical stimulation with PMA and ionomycin, in the presence of a Golgi inhibitor (Brefeldin A). A substantial portion of aged T cells (approxiamately 26% of CD4^+^ T cells and over 80% of CD8^+^ T cells) produced IFNγ, significantly more than the portion of young T cells (figure 4A, B). Moreover, while the percentage of young T cells producing GzB was negligible, aged T cells showed a significant increase in GzB production, particularly among CD8^+^ T cells (figure 4C,D). Consistent with these findings, about 10% of T cells derived from the spleens of aged mice expressed CD107, a marker for degranulation and secretory activity, compared to only 0.6% of young T cells (figure 4E,F). CD107 expression levels were similar in aged CD4^+^ and CD8^+^ T cells (supplemental figure 2A).

**Figure 4:**
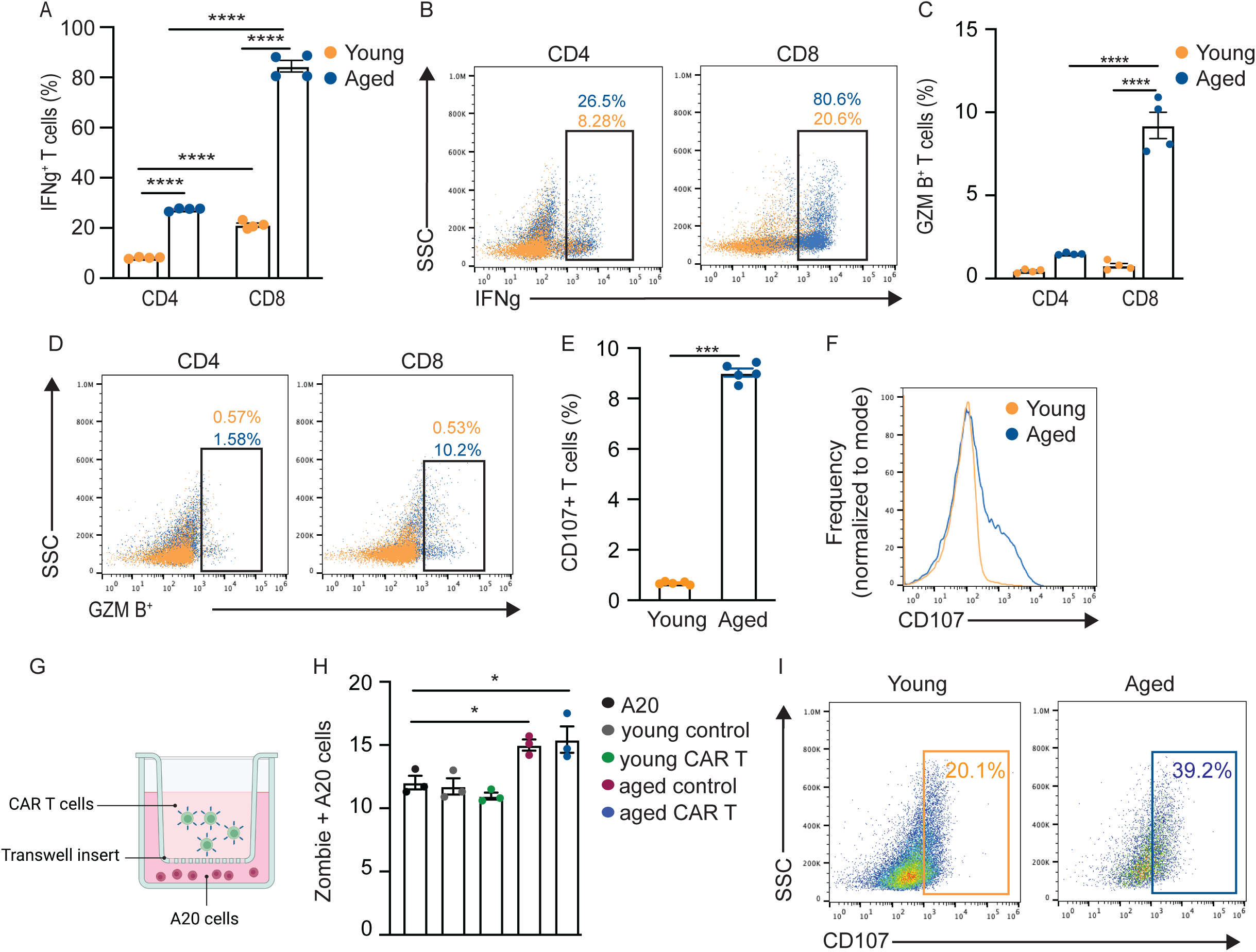
Aged T cells eliminate target cells through the continuous secretion of GzB and IFNγ. Young and aged T cells were subjected to a brief chemical stimulation with PMA and ionomycin in the presence of a Golgi inhibitor (BrefeldinA) for 5 hours. Production of IFNγ (A,B) and GzB (C,D) was quantified by flow cytometry. Each point represents data collected from a single young (n=4) or aged (n=4) mouse. (E,F) Analysis of CD107 expression on freshly isolated young and aged T cells. Each point represents data collected from a single young (n=5) or aged (n=5) mouse (G) Schematic showing the setup of a Transwell co-culture assay. (H) Quantitation of A20 killing by young and aged CAR T cells using the Transwell co-culture system. T cells used for CAR T cell generation were pooled from young (n=3) and aged (n=5) mice. Single dots represent technical replicates. (I) Representative plots showing CD107 expression in young vs. aged CAR T cells. Bar graphs represent mean ± SEM. (*p<0.05, ****p<0.0001); one-way ANOVA with Tukey’s multiple comparisons test (A,C,H), or unpaired Student’s t test (E).

These findings are consistent with previous studies, suggesting that aged T cells exist in a constant state of degranulation^23^, which could explain why specific target recognition was not required for cytotoxic activity. To test this hypothesis, CAR T cells and control T cells derived from young and aged donor mice were cultured with A20 target cells using a Transwell setup, where effector and target cells were placed in separate compartments (figure 4G). Aged, but not young, T cells induced target cell death, even in the absence of physical contact (figure 4H). Consistently, CD107 expression was twice as high in aged compared to young CAR T cells (figure 4I). These results suggest that aged T cells kill their targets through continuous degranulation and secretion of cytotoxic cytokines.

### Young T cells acquire a proinflammatory phenotype when transferred into an aged host

Our previous studies demonstrated that exposure to the hemolytic microenvironment in aged spleens induces multiple aging phenotypes in young T cells, including reduced proliferation, and upregulation of CD39^21^. To test whether exposure to the aged microenvironment in the spleen contributed to the continuous degranulation and cytotoxicity observed in aged T cells, TdTomato^+^ T cells were isolated from young, transgenic donors and transferred to young or aged C57Bl/6 wild-type recipients. The mice were sacrificed after 3 weeks, and spleens were collected for analyzing the secretory phenotype of transferred, TdTomato^+^ T cells (figure 5A).

**Figure 5:**
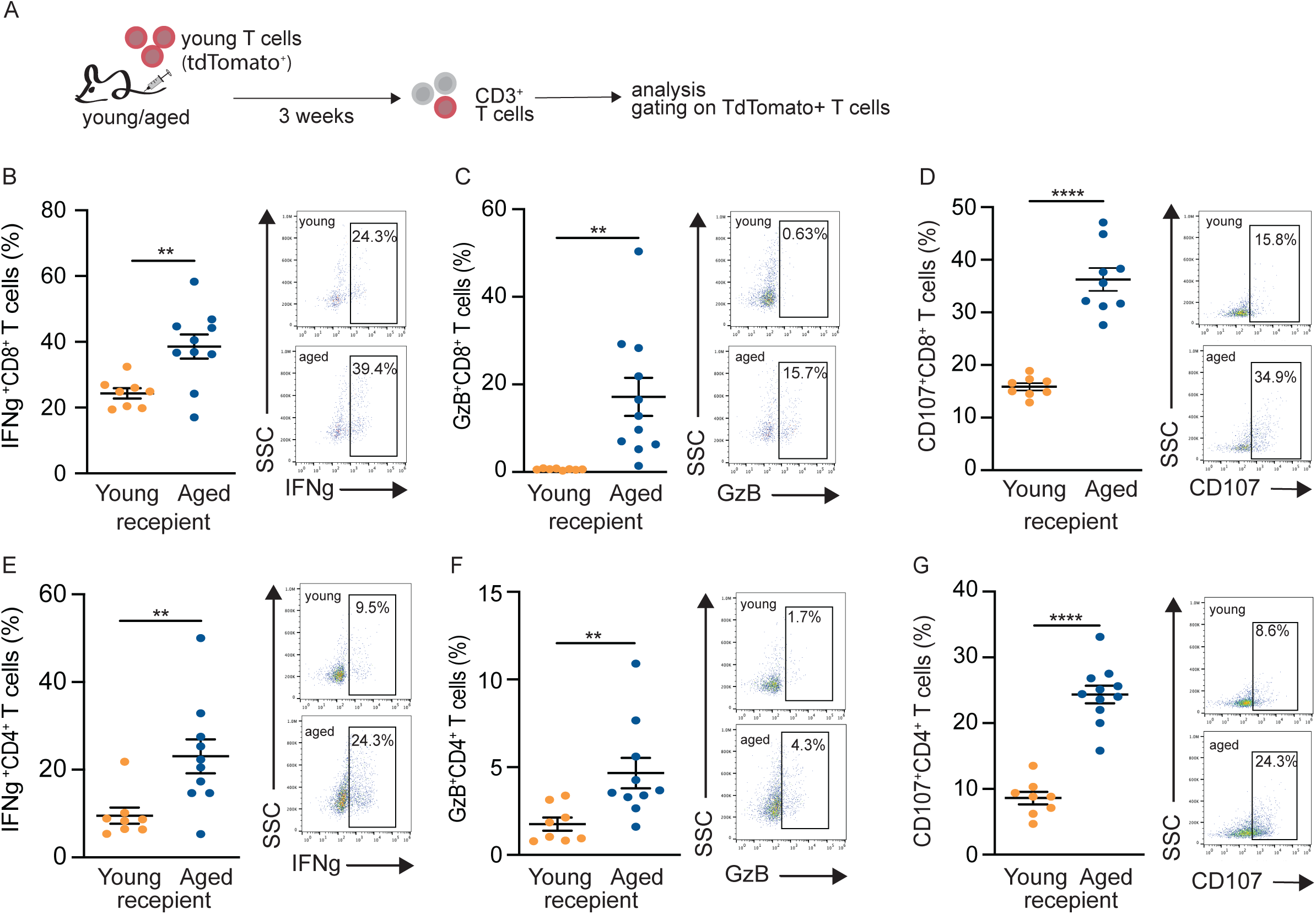
Young T cells acquire a proinflammatory phenotype when transferred into an aged host. (A) Experimental design. Young T cells from transgenic mice constitutively expressing TdTomato were transferred into young or aged C57Bl/6 wild-type recipients. After 3 weeks, recipient mice were sacrificed, and CD3^+^ T cells were purified from the spleen, and analyzed by flow cytometry, gating on TdTomato^+^ T cells. Production of IFNγ and GzB was analyzed after a short ex vivo stimulation with PMA and ionomycin in the presence of a Golgi inhibitor (BrefeldinA). CD107 expression was quantified on freshly isolated T cells. CD8^+^ T cells (B-D), and CD4^+^ T cells (E-G) were analyzed separately. Error bars represent mean ± SEM. Each dot represents data collected from a single young (n=8) or aged (n=11) recipient mouse (**p<0.01, ****p<0.0001); Student’s t-test.

Similar to aged CD8^+^ T cells, young CD8^+^ T cells isolated from aged spleens exhibited increased IFNγ (figure 5B) and GzB (figure 5C) production, along with higher surface expression of CD107(figure 5D), indicating elevated degranulation. Interestingly, young CD4^+^ T cells also showed increased cytokine production and cytotoxicity after exposure to an aged microenvironment (figure 5E-G), though to a lesser extent than CD8^+^ T cells. Previous studies identified elevation of GzB secretion and degranulation in aged T cells^23^. Furthermore, a cytotoxic CD4^+^ T cell subpopulation was identified that accumulates with aging^24,25^. Our findings suggest these phenotypes are driven, at least in part, by the aged microenvironment.

### The efficacy of CAR T cell-mediated immunotherapy is reduced in aged compared to young tumor-bearing mice

To investigate whether host age affects the anti-tumoricidal response of CAR T cells in immune-competent mice, we utilized the syngeneic ZsGreen^+^ ML21 B cell leukemia model. In this system, B cell leukemia was induced in situ through the infusion of bone marrow cells transduced with multiple oncogenes driven by a tetracycline-responsive promoter^15^. After confirming that ML21 leukemia cells express CD19, the target antigen for our CAR T cells (supplemental figure 3A), we transfused ML21 cells into young and aged mice pre-treated with cyclophosphamide. Doxycycline was administered in drinking water throughout the experiment to induce transformation (supplemental figure 3B). Peripheral blood analysis, starting on day 8, showed a gradual accumulation of leukemia cells (ZsGreen^+^). By day 16, approximately 60% of lymphocytes in both young and aged mice were leukemia cells (supplemental figure 3C). Notably, in this first experiment, two mice did not develop leukemia, suggesting some variability in disease induction (supplemental figure 3C). Based on these data, we selected day 10 for CAR T cell administration in following experiments, as this timepoint represented a linear phase of disease progression with comparable ML21 burdens across individual mice (supplemental figure 3D). To accurately assess treatment efficacy without ongoing disease induction, doxycycline was discontinued at the time of therapy initiation (supplemental figure 3D).

While disease progression was more pronounced in mice treated with control T cells, by day 15, we observed a spontaneous decrease in ML21 frequency in both groups (supplemental figure 3E). To mitigate this spontaneous disease resolution, we modified our protocol to maintain doxycycline administration throughout the experiment (figure 6A). Using this optimized protocol, young tumor-bearing mice were treated with either control or CAR T cells. ML21 levels in the blood continued to rise until day 15, after which we observed a significant reduction in leukemia burden in CAR T cell-treated mice but not in controls (figure 6B). Thus, despite ongoing doxycycline administration, we successfully established an in vivo protocol to evaluate CD19 CAR T cell therapy in C57Bl/6 immunocompetent mice.

**Figure 6:**
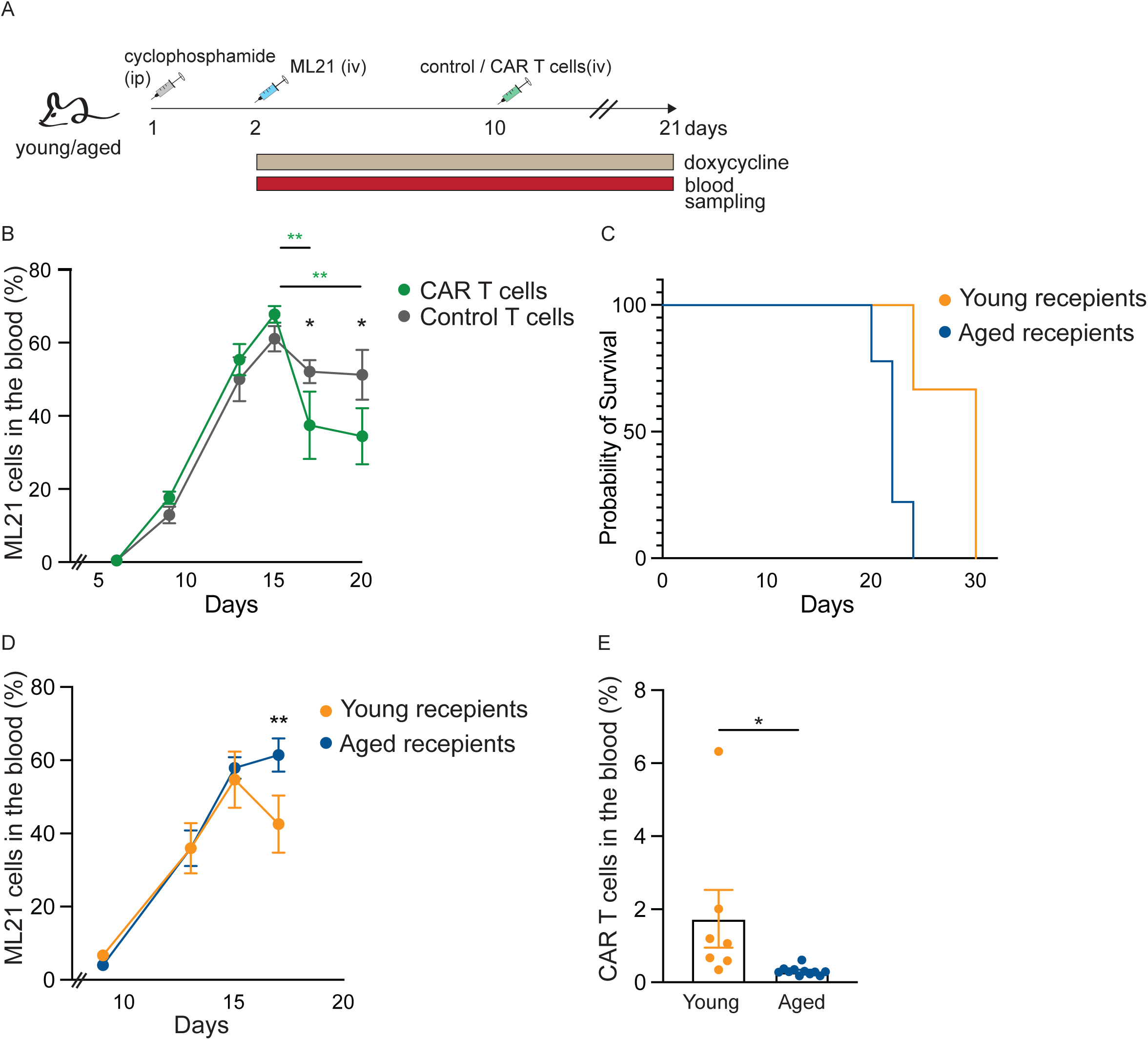
The efficacy of CAR T cell-mediated immunotherapy is reduced in aged compared to young tumor bearing mice. (A) Experimental design. Young and aged mice were injected intraperitoneally (i.p.) with cyclophosphamide, followed by transfusion with genetically modified bone marrow cells. Doxycycline was administered in drinking water to induce transformation. On day 10, the mice received an i.v. injection of either control or CAR T cells. Blood samples were collected via tail vein punctures. (B) ML21 leukemia was established in 8 young mice, followed by transfusion of either control (n=4) or CAR (n=4) T cells. Percentage of ML21 from total lymphocytes was determined by quantifying ZsGreen expression. (C-E) Young (n=6) and aged (n=10) mice were inoculated with ML21 leukemia, and treated with young CAR T cells. (C) Kaplan-Meier survival curve. (D) Percentage of ML21 from total lymphocytes was determined by quantifying ZsGreen expression. (E) Percentage of CAR T cells of total lymphocytes was determined using the Myc tag incorporated into the CAR. Data points represent mean ± SEM. (*p<0.05, **p<0.01); two-way ANOVA with Tukey’s post-hoc analysis (B, D), and unpaired Student’s t-test (E).

To assess the impact of an aged microenvironment on CAR T cell efficacy in vivo, ML21 leukemia was established in six young and ten aged mice. All mice received young CAR T cells on day 10, following the protocol outlined in Figure 6A. Survival was significantly better in young mice treated with CAR T cells, compared to aged CAR T cell-treated mice (figure 6C). Moreover, while young mice exhibited a marked reduction in ML21 levels following CAR T cell treatment, aged tumor-bearing mice did not respond and retained high ML21 burdens (figure 6D). Consistent with the lower efficacy observed in aged mice, and despite transfusing similar numbers of CAR T cells to all mice, quantification of CAR T cells in peripheral blood revealed significantly reduced CAR T cell persistence in aged hosts (figure 6E).

Our findings highlight that the age of a tumor-bearing mouse significantly influences the efficacy of CAR T cell immunotherapy at multiple levels. Aging reduces CAR T cell production yield, and alters CAR cell function. While exposure to an aged microenvironment enhances T cell and CAR T cell cytotoxicity, this cytotoxic effect is non-specific, and CAR T cells in aged hosts exhibit reduced persistence and impaired tumor clearance in vivo. This aligns with their altered subpopulation composition, which favors an effector memory phenotype over the central memory phenotype seen in younger CAR T cells—an attribute known to impact persistence^26^.

## Discussion

Aging is associated with a progressive decline in both innate and adaptive immune functions, contributing to increased susceptibility to infections, reduced vaccine efficacy, and a higher risk of malignant and inflammatory diseases^27–29^. Among immune cell populations, T cells are particularly affected by aging. Thymic involution leads to diminished naïve T cell production, while cumulative cellular and molecular defects impair T cell activation, proliferation, and memory formation^1^. Additionally, age-related changes in antigen presentation and stromal cell composition within the lymphoid microenvironment further compromise T cell functionality^30^. Given the essential role of T cells in cancer immunotherapy, this study aimed to elucidate how these age-associated changes impact the efficacy of CAR T cell therapy.

Our findings demonstrate that aging influences CAR T cell therapy at multiple stages, starting from production of the transduced cells. T cells derived from aged mice exhibit reduced transduction efficiency, and resulted in an altered CAR T cell product composition. Aged CAR T cells contain a higher proportion of CD4^+^ T cells with an effector memory phenotype, as opposed to the dominance of the CD8^+^ central memory phenotype observed in younger CAR T cells. Given that CD8⁺ central memory cells are known for their superior expansion and long-term persistence, while CD4⁺ effector memory cells have limited proliferative capacity and persistence, this age-related change in subpopulation composition may negatively impact CAR T cell durability and therapeutic efficacy^31,32^. Thus, a patient’s immunological age may influence the characteristics of their CAR T cell product and, consequently, the therapy’s effectiveness.

Previous studies have reported conflicting results regarding the cytotoxicity of young versus aged CAR T cells^11–13^. Our studies show that while aged CAR T cells exhibit reduced killing efficacy, they display non specific cytotoxicity, as evidenced by increased degranulation and GzB secretion independent of target recognition. We further show that this phenotype is not intrinsic to aged T cells but is induced by exposure to the aged systemic milieu. Accordingly, even young T cells aqcuired this phenotype when transferred into aged hosts. Non-specific cytotoxicity could harm healthy tissues, and is in line with reports of elevated risk for cytokine release storm (CRS) in aged patients undergoing CAR T cell therapy. Importantly, the demonstrated differences in outcomes of CAR T cell therapy between young and aged tumor-bearing mice likely stem from a combination of factors, including the accumulation of suppressor cells in aged hosts, and the overall inflammatory milieu that may promote activation-induced cell death (AICD)^27–29^. Our previous work identified the hemolytic microenvironment of the aged spleen—characterized by iron and heme accumulation—as a driver of T cell dysfunction, including the upregulation of exhaustion markers and reduced proliferative capacity^21^. We further demonstrated that aged T cells exhibit resistance to ferroptosis, an iron- and oxidative stress-induced form of cell death, by limiting iron uptake^21^. While this adaptation enhances survival in a stressful microenvironment, it comes at the cost of impaired proliferation, which relies on iron-dependent processes such as mitochondrial biogenesis and DNA synthesis^33,34^. Thus, the aged microenvironment may impair CAR T cell persistence in aged hosts by diminishing their proliferative capacity.

In this study we used a second generation CAR containing a co-stimulatory domain derived from CD28, known to promote an effector memory phenotype. It is possible that other CAR designs could improve CAR T cells function and persistence in the aged milieu. For example incorporating other costimulatory domains like 4-1BB were shown to enhance T cell persistence^35^. Other approaches to optimize CAR T cell fitness, like the co-expression of key metabolic enzymes^36^, the elimination of inhibitory signals^37^ could further help improve the long-term efficacy of CAR T cells in older patients.

Our findings, if proven relevant to human patients, will have important implications for CAR T cell therapy, particularly in the context of aging and “off-the-shelf” allogeneic CAR T therapies^38^. Universal donor-derived CAR T cells are an attractive solution to overcome the logistical challenges of autologous CAR manufacturing. However, our results suggest that the efficacy of such treatments may be influenced by the recipient’s immune status and the aged lymphoid microenvironment. Understanding and addressing these age-related barriers will be critical in optimizing CAR T cell therapies for older patients.

## Acknowledgements

The authors thank Prof. Ziv Shulman for providing mouse models. Dr. Shelley Schwarzbaum for her assistance in editing the manuscript. Viktoria Zlobin and Dr. Amit Avrahami from the Technion Preclinical Authority for maintaining our aged mouse colony and help with in vivo studies. Dr. Aviv Lutati and Yousef Mansour from the Technion Life Science and Engineering Infrastructure Center for their help with cell sorting. The research was funded by grants given to N.R.-H. by the Israeli Cancer Association (ICA) and the Israel Cancer Research Fund (ICRF).

## Supplementary Figure Legends

**Supplemental Figure 1:**

(A) Schematic of the PBullet plasmid encoding a CD19-targeting CAR with the GFP reporter.

(B) Analysis of A20 killing after coculture with young control and CAR T cells, at varying T: E ratios. A20 cell killing was evaluated by flow cytometry using PI staining. (C) CD69 expression on CD3⁺ T cells from young and aged mice after weak activation. (D-E) CAR transduction efficiency in young and aged T cells stimulated with varying doses of anti-CD28 (D) or anti-CD3 (E). (F) Analysis of CD69 expression on CD3⁺ T cells from young and aged mice following strong activation. Bar graphs represent mean ± SEM. Data are pooled from 3 young mice; each dot represents a technical replicate (****p<0.0001); one-way ANOVA with Tukey’s multiple comparisons test.

**Supplemental Figure 2:**

(A) CD107 expression on freshly isolated young (n=5) and aged (n=5) T cells. Analyzed by flow cytometry.

**Supplemental Figure 3:**

(A) CD19 expression on ZsGreen⁺ ML21 cells analyzed by flow cytometry. (B) Experimental design. Young and aged mice were injected intraperitoneally (i.p.) with cyclophosphamide, followed by i.v. transfusion of ML21 leukemia cells. Doxycycline was administered in drinking water. Blood samples were collected via tail vein punctures. (C) ML21 leukemia was established in young (n=5) and aged (n=7) mice. ML21 percentage out of total lymphocytes was determined by quantifying ZsGreen expression. (D) Experimental design. ML21 leukemia was established in 10 young mice, followed by transfusion of either control (n=5 recipients) or CAR (n=5 recipients) T cells. Doxycycline was withdrawn at the time of treatment. (E) Analysis of ML21 percentage of total lymphocytes in the experiment described in D, determined by quantifying ZsGreen expression. (***p<0.001); two-way ANOVA Tukey’s post-hoc analysis.

