## Supplemental Figures for "An Aged Microenvironment Increases CAR T Cell Cytotoxicity but Impairs Therapeutic Efficacy"

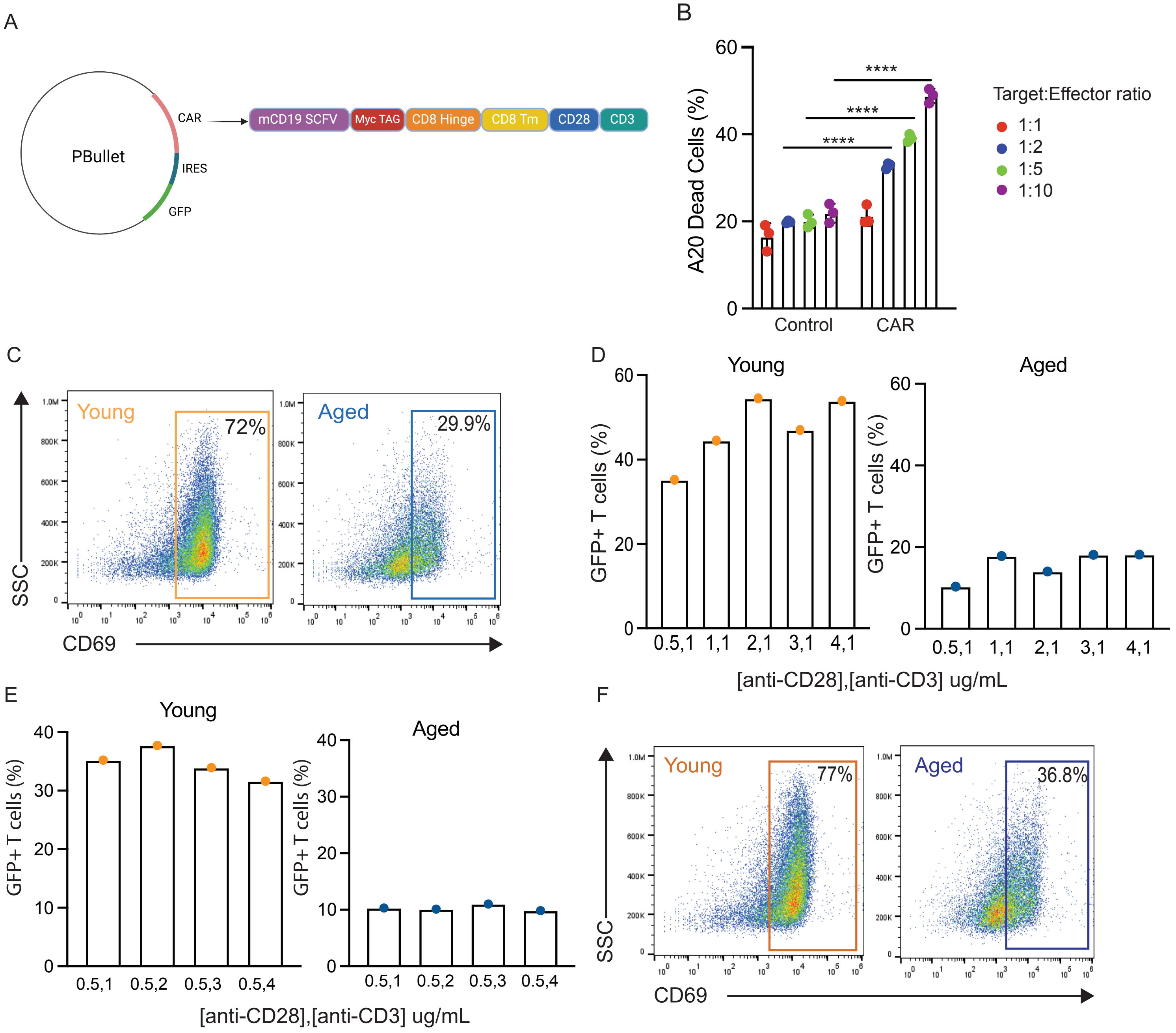


**Supplemental Figure 1:**
(A) Schematic of the PBullet plasmid encoding a CD19-targeting CAR with the GFP reporter. (B) Analysis of A20 killing after coculture with young control and CAR T cells, at varying T: E ratios. A20 cell killing was evaluated by flow cytometry using PI staining. (C) CD69 expression on $CD3^{+}$ T cells from young and aged mice after weak activation. (D-E) CAR transduction efficiency in young and aged T cells stimulated with varying doses of anti-CD28 (D) or anti-CD3 (E). (F) Analysis of CD69 expression on $CD3^{+}$ T cells from young and aged mice following strong activation. Bar graphs represent mean ± SEM. Data are pooled from 3 young mice; each dot represents a technical replicate (****$p<0.0001$); one-way ANOVA with Tukey's multiple comparisons test.

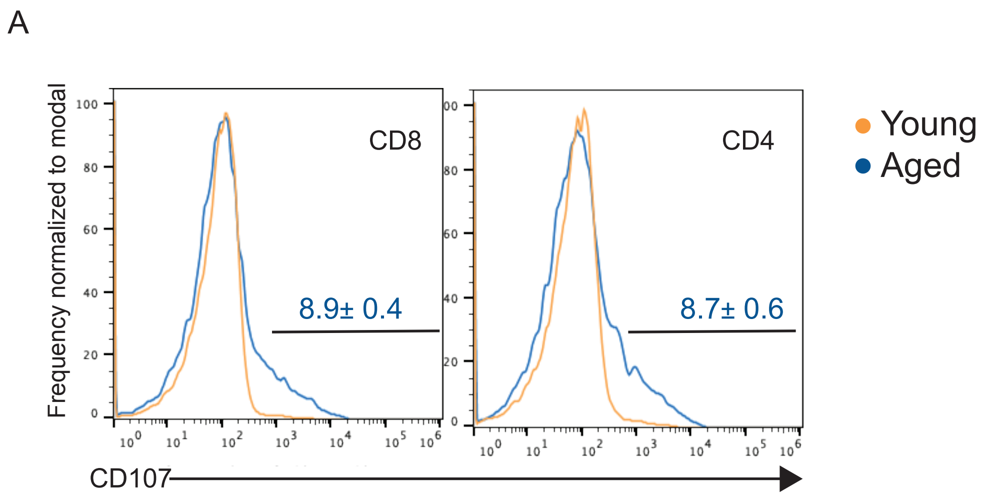


**Supplemental Figure 2:**
(A) CD107 expression on freshly isolated young (n=5) and aged (n=5) T cells. Analyzed by flow cytometry.

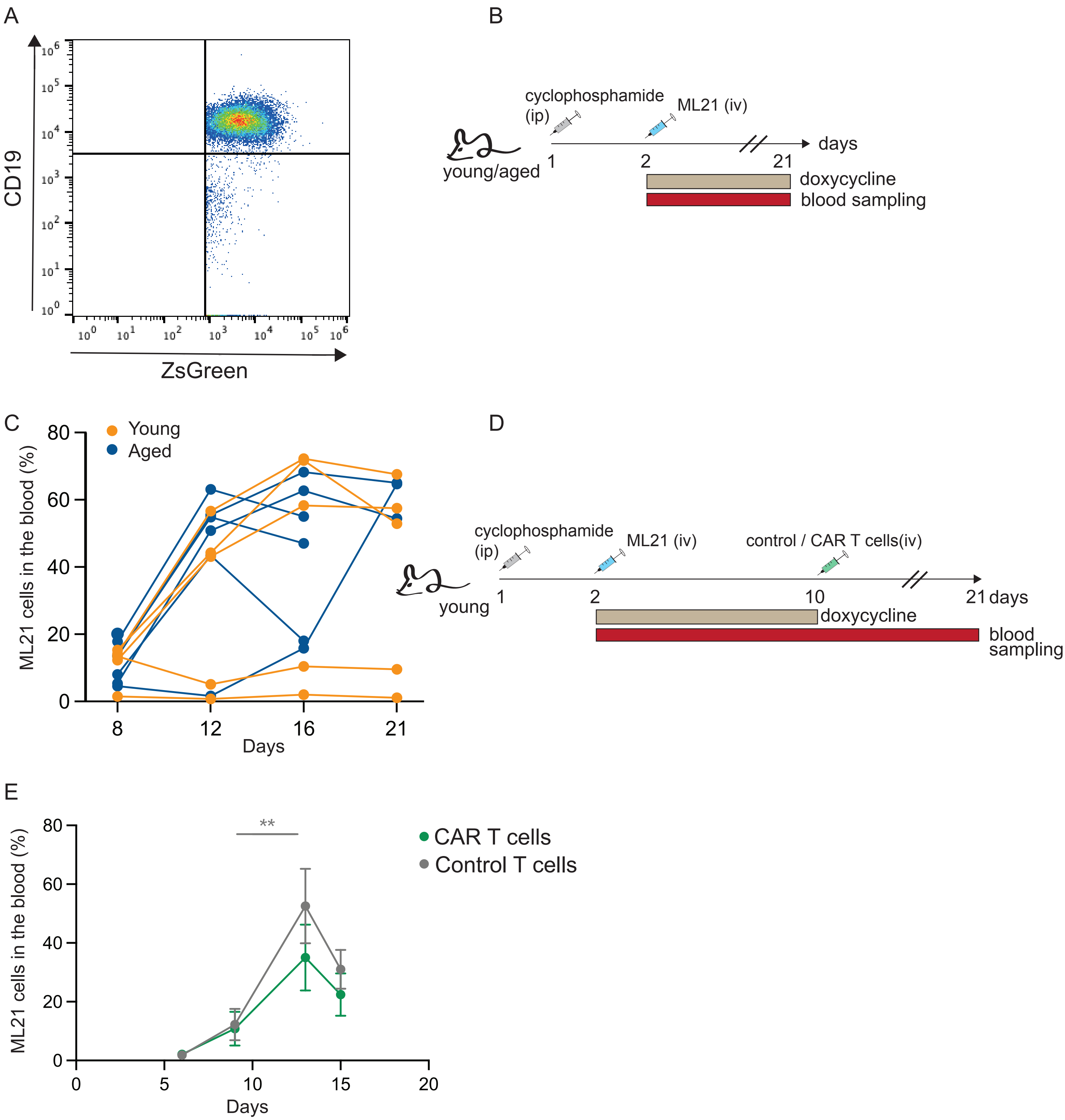


**Supplemental Figure 3:**
(A) CD19 expression on ZsGreen$^{+}$ ML21 cells analyzed by flow cytometry. (B) Experimental design. Young and aged mice were injected intraperitoneally (i.p.) with cyclophosphamide, followed by i.v. transfusion of ML21 leukemia cells. Doxycycline was administered in drinking water. Blood samples were collected via tail vein punctures. (C) ML21 leukemia was established in young (n=5) and aged (n=7) mice. ML21 percentage out of total lymphocytes was determined by quantifying ZsGreen expression. (D) Experimental design. ML21 leukemia was established in 10 young mice, followed by transfusion of either control (n=5 recipients) or CAR (n=5 recipients) T cells. Doxycycline was withdrawn at the time of treatment. (E) Analysis of ML21 percentage of total lymphocytes in the experiment described in D, determined by quantifying ZsGreen expression. (***$p<0.001$); two-way ANOVA Tukey's post-hoc analysis.
